# A molecular mechanism for salt stress-induced microtubule array formation in Arabidopsis

**DOI:** 10.1101/431031

**Authors:** Christopher Kesten, Arndt Wallmann, René Schneider, Heather E. McFarlane, Anne Diehl, Ghazanfar Abbas Khan, Barth-Jan van Rossum, Edwin Lampugnani, Nils Cremer, Peter Schmieder, Kristina L. Ford, Florian Seiter, Joshua L. Heazlewood, Clara Sanchez-Rodriguez, Hartmut Oschkinat, Staffan Persson

## Abstract

Microtubules are filamentous structures necessary for cell division, motility and morphology, with dynamics critically regulated by microtubule-associated proteins (MAPs). We outline the molecular mechanism by which the MAP, COMPANION OF CELLULOSE SYNTHASE1 (CC1), controls microtubule bundling and dynamics to sustain plant growth under salt stress. CC1 contains an intrinsically disordered N-terminus that links microtubules at evenly distributed distances through four conserved hydrophobic regions. NMR analyses revealed that two neighboring residues in the first hydrophobic binding motif are crucial for the microtubule interaction, which we confirmed through live cell analyses. The microtubule-binding mechanism of CC1 is remarkably similar to that of the prominent neuropathology-related protein Tau, indicating evolutionary convergence of MAP functions across animal and plant cells.

## Introduction

Microtubules are tubular structures essential to morphogenesis, division and motility in eukaryotic cells ^1^. While animal cells typically contain a centrosome with radiating microtubules toward the cell periphery, growing plant cells arrange their microtubules along the cell cortex ^2^. A major function of the cortical microtubules in plant cells is to direct the synthesis of cellulose, a fundamental component of the cell wall essential to plant morphology ^3^. Cellulose is produced at the plasma membrane by Cellulose Synthase (CESA) protein complexes (CSCs; ^4^) that display catalytically-driven motility along the membrane ^3^. The recently described microtubule-associated protein (MAP), Companion of Cellulose Synthase^1^ (CC1), is an integral component of the CSC and sustains cellulose synthesis by promoting the formation of a stress-tolerant microtubule array during salt stress ^5^. As cellulose synthesis is key for plant growth, engineering of plants to better produce cellulose is of utmost importance to agriculture. Indeed, understanding the molecular mechanism by which CC1 controls cellulose synthesis may bear opportunities to improve cultivation on salt-affected lands.

The microtubule network is highly dynamic, and its state is influenced by the action of MAPs. The mammalian Tau/MAP2/MAP4 family represents the most investigated MAP set, primarily due to Tau’s importance in the pathology of neurodegenerative diseases ^6–8^. *In vitro*, Tau promotes polymerization and bundling of microtubules, and diffuses along the microtubule lattice ^9–11^. In the brain, Tau is predominantly located at the axons of neurons, where it contributes to the microtubule organization that drives neurite outgrowth ^12,13^. In disease, Tau self-aggregates into neurofibrillary tangles that might trigger neurodegeneration ^14^. Intriguingly, no clear homologs of the Tau/MAP2/MAP4 family have been identified in plants ^15,16^. Because the full scope of Tau’s biological role remains elusive, identification of Tau-related proteins outside the animal Kingdom would benefit our understanding of how this class of MAPs functions.

### The N-terminus of CC1 Links Microtubules at Evenly Distributed Distances and Bundles Microtubules

The cytosolic N-terminal part of CC1 (residues 1-120, CC1ΔC223) binds to microtubules and restores microtubule re-assembly, cellulose synthesis and wild-type growth of *cc1cc2* (null-mutation in CC1 and its closest homolog CC2) seedlings on high levels of salt ^5^. These data indicate that CC1ΔC223 is critical to CC1’s function during stress, and we therefore set out to investigate the molecular details of how it interacts with microtubules. We cross-linked 6xHis-tagged CC1ΔC223 with α-β-tubulin dimers using 1-ethyl-3-(3-dimethylaminopropyl) carbodiimide hydrochloride (EDC) ^17^, which led to di- and multimeric protein products (Fig. 1A and Fig. S1A). LC/MS/MS analysis revealed five well-defined covalent bonds between CC1ΔC223 and α- or β-tubulin (Fig. 1B). We consistently detected four peptides of CC1ΔC223 cross-linked to β-tubulin (K^40^ to E^111^, K^94^ to E^111^, K^96^ to E^111^ and K^96^ to E^158^; letters and numbers indicate amino acids in CC1ΔC223 and β-tubulin, respectively; Fig. 1, B to C; Table S1). Notably, the three sequentially distant K^40^ and K^94/96^ of CC1ΔC223 cross-linked to the same residue on β-tubulin (E^111^). This suggests that two CC1 regions might bind the same sites on two different β-tubulin molecules, which is corroborated by the multimeric protein products in the SDS page. The cross-linked position on α-tubulin is close to the hydrophobic interface between tubulin heterodimers, a site that is frequently occupied by agents that directly regulate microtubule formation such as vinblastine, the stathmin-like domain (SLD) of RB3, and also by Tau ^18–20^ (Figs. S1B and C).

**Figure 1.**
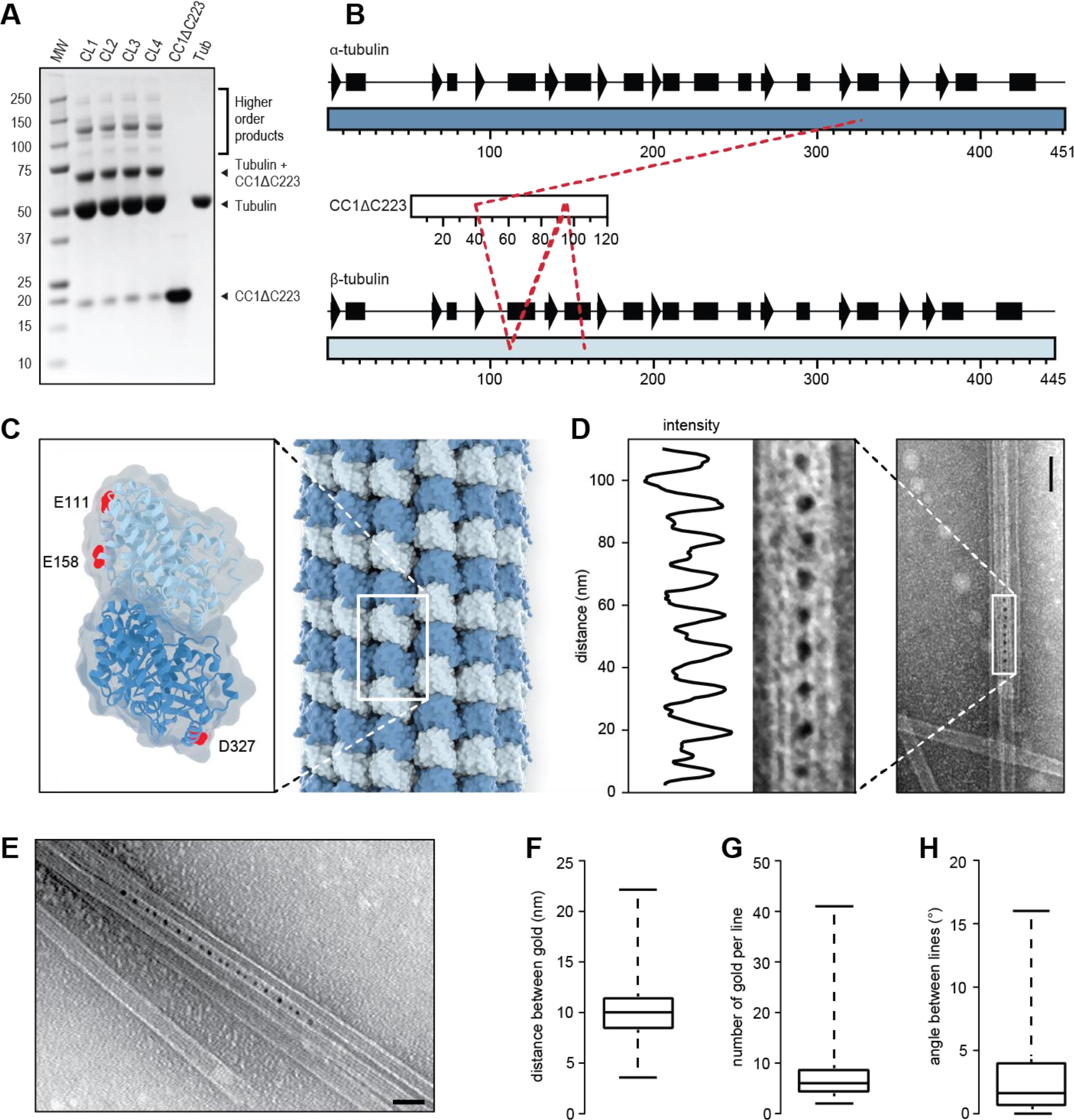
The N-terminus of CC1 binds sites on both α- and β-Tubulin and cross-links microtubules. **A.** SDS-Page of EDC-induced cross-linking of 6xHis-CC1ΔC223 (16 kDa) and tubulin dimers (2 × 55 kDa). Arrowheads depict position of relevant protein bands. MW = molecular weight marker, CL1-4 = cross-linking reaction 1-4. Higher order cross-linking products represent cross-links between e.g. tubulin + 2 × CC1ΔC223 (87 kDa), tubulin dimers (110 kDa), tubulin dimers + CC1ΔC223 (126 kDa), tubulin dimers + 2 × CC1ΔC223 (142 kDa). **B.** Schematic views of the secondary structures of α- and β-tubulin, and the CC1ΔC223 sequence. Dashed lines depict detected cross-linking positions of CC1ΔC223 and α- or β- tubulin. **C.** Projection of detected cross-links onto an α/β-tubulin dimer. Dark blue = α-tubulin; Light blue = β-tubulin; Sites for cross-linked amino acids are marked in red. **D.** Representative TEM image of CC1ΔC223 distribution along negatively-stained, taxol-stabilized microtubules polymerized in the presence of 6xHis-CC1ΔC223. CC1ΔC223 protein is visualized by a 5 nm gold-conjugated Ni-NTA tag that recognizes 6xHis-tagged proteins. A transect was taken along rows of gold particles, and dips in the light intensity along the transect correspond to gold particle centers. Note the even distribution of the electron-dense gold particles in between neighboring microtubules. Scale bar = 50 nm. **E.** CC1ΔC223 distribution along negatively-stained, taxol-stabilized microtubules polymerized in the presence of 6xHis-CC1ΔC223. CC1ΔC223 can form a zipper-like pattern that links microtubules. Scale bar = 100 nm. **F.** Quantification of the distance between individual gold particles as shown in **D** and **E** (box plot: Center lines show the medians; box limits indicate the 25th and 75th percentiles; whiskers extend to the minimum and maximum). **G** and **H.**Quantification of number of gold labels per row (**G**) and the angle between adjacent gold-labeled rows (**H**) from images as those in **D** and **E** (box plots: Centerlines show the medians; box limits indicate the 25th and 75th percentiles; whiskers extend to the minimum and maximum).

To further investigate how CC1ΔC223 binds microtubules, we co-polymerized tubulin in the presence of CC1ΔC223. We then labeled CC1ΔC223 using 5 nm gold-conjugates that recognize the His-tag ^21^ and monitored the formed microtubules and gold distribution *via* transmission electron microscopy (TEM). Gold labeling only occurred at closely aligned microtubules with very small inter-microtubule distances (Fig. 1, D and E; Fig. S1D), and occurred as evenly distributed distances in straight rows along interphases of two neighboring microtubules (Fig. 1, D and E). The gold particles were typically spaced by 10 nm (Fig. 1F; 10.0 nm ± 2.4 nm; mean +/− SD; three independent experiments; N = 1785 labels). The number of gold-labels in a given row ranged between two and 41 labels (Fig. 1G; 8 ± 5 labels; mean +/− SD; three independent replicates; N = 274 rows), making each row about 80 nm in length. We also observed multiple gold-labeled rows on one microtubule when in close proximity to several other microtubules (Fig. 1E). The angles between gold-labeled rows were small (Fig. 1H; 2.8° ± 3°; mean +/− SD; three independent replicates; N = 98 rows), highlighting that the labeling did not shift between neighboring protofilaments on the same microtubule. These data indicate that CC1ΔC223 promotes microtubule bundling. Indeed, increasing levels of CC1ΔC223 correlated with increased microtubule bundling in TEM experiments (Fig. 2, A to C; Fig. S2, A and B).

**Figure 2.**
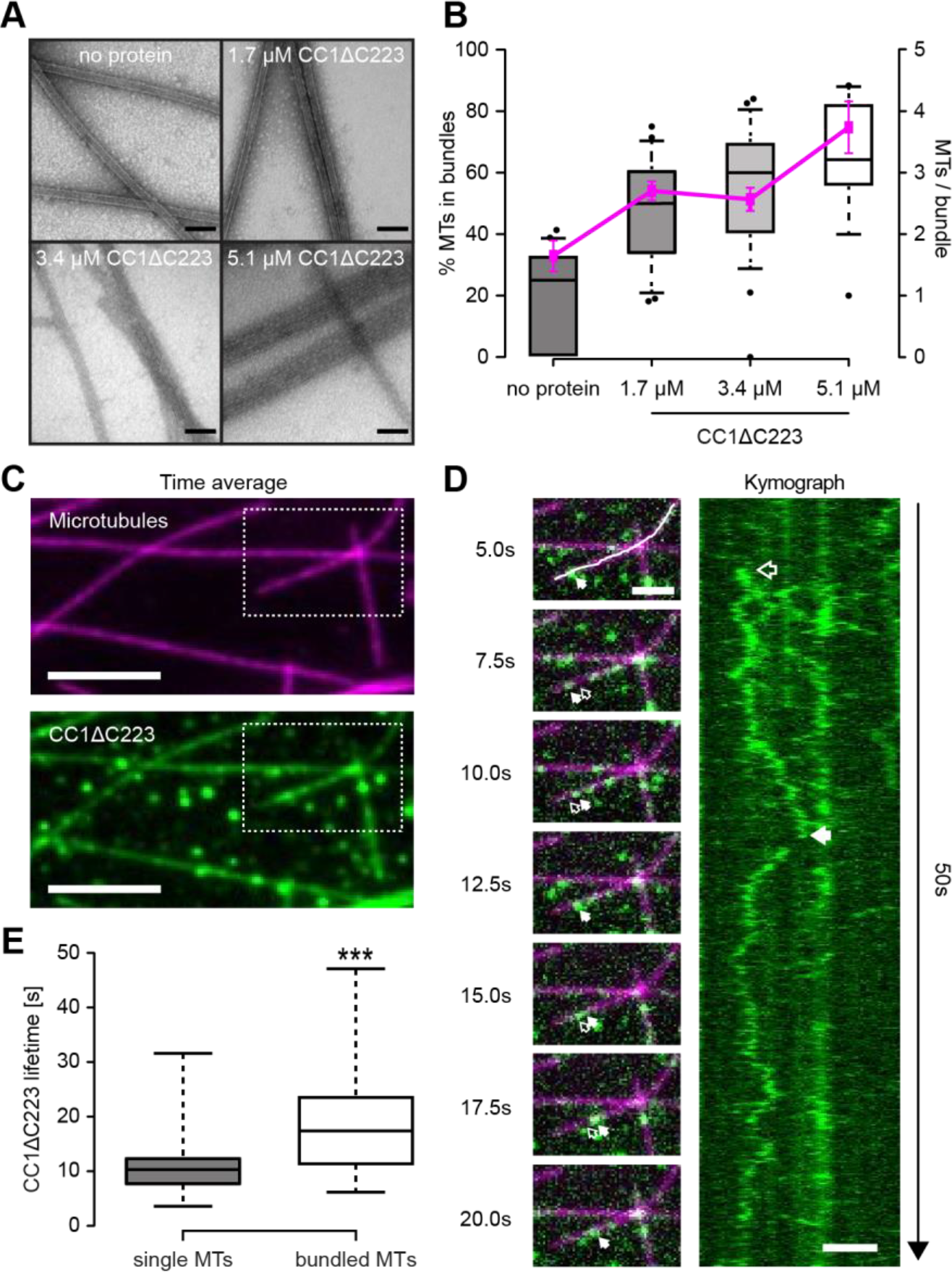
The N-terminus of CC1 induces microtubule bundling and can diffuse along the microtubule lattice. **A.** Transmission electron microscopy (TEM) of negatively-stained taxol-stabilized microtubules after addition of increasing levels of 6xHis-CC1ΔC223 during microtubule polymerization. Note that it is very difficult to discern individual microtubules in the microtubule bundles after addition of ~ 3 µM of CC1ΔC223. Scale bars = 100 nm. **B.** Quantification of the proportion of microtubules in bundles (left y-axis, box plots: Center lines show the medians; box limits indicate the 25th and 75th percentiles; whiskers extend to the 10th and 90th percentiles, outliers are represented by dots) and number of microtubules/bundle (right y-axis, magenta line: mean +/− SEM) with increasing concentration of H6xis-CC1ΔC223 (quantified from images such as those in A). **C.** CF488A-labeled 6xHis-CC1ΔC223 proteins (green) associated with surface-bound microtubules (magenta) *in vitro*. Scale bar = 5 µm. **D.** Time-series images (left panel) of CF488-labeled 6xHis-CC1ΔC223 (green) diffusing along microtubules (magenta). Filled arrow = position in current frame, empty arrow = position in previous frame. Scale bar = 2 µm. Representative kymograph (right panel) along solid line in left panel (top) showing diffusion of 6xHis-CC1ΔC223 foci. Scale bar = 2 µm. **E.** 6xHis-CC1ΔC223 lifetime on single versus bundled microtubules (box plots: Center lines show the medians; box limits indicate the 25th and 75th percentiles; whiskers extend to the minimum and maximum), n = 60 single and 37 bundled microtubules, *** p-value < 0.001, Welch’s unpaired *t*-test).

### The N-terminus of CC1 can Diffuse Along the Microtubule Lattice

As our TEM experiments only provide static information on the interactions between CC1ΔC223 and microtubules, we labelled the sole sulfhydryl group (C^116^) in CC1ΔC223 with the green fluorescent dye CF488A-maleimide (Fig. S2C) and performed rhodamine-labeled microtubule interaction assays ^22^. Using total internal reflection fluorescence microscopy, we observed most of the CF488A-labeled CC1ΔC223 proteins as fluorescent foci associated with microtubules (Fig. 2C; Movie S1). CF488A-labeled CC1ΔC223 diffused bidirectionally along the rhodamine-labeled microtubules and occurred on both single and bundled microtubules (Fig. 2D). The mean square displacement (0.076 ± 0.005 µm^2^/s; mean ± S.D.; *N* = 50 molecules) of fluorescent foci exhibited a linear relationship with time (Fig. S2, E and F), indicating free diffusion. In accordance with the results above, CF488A-labeled foci occupied bundled microtubules for a longer time than single microtubules (Fig. 2E; fig. S2, D and E). These data are reminiscent to that of Tau, which promotes microtubule-bundling and polymerization, and also moves along microtubules *in vitro* with comparable diffusion coefficients (0.142 - 0.292 µm^2^/s; 10).

### The N-terminus of CC1 is Intrinsically Unstructured and Engages with Microtubules through Four Hydrophobic Motifs

To understand how CC1ΔC223 engages with microtubules, we assessed its structural features using solution state NMR, circular dichroism spectroscopy (CD) and analytical ultracentrifugation (AUC). The 2D ^1^H-^15^N-heteronuclear single quantum coherence spectrum of ^15^N-labeled CC1ΔC223 showed narrow signals and poor chemical shift dispersion in the ^1^H dimension, which is characteristic for intrinsically disordered proteins (Fig. 3A). For the sequence-specific assignment, we used a combination of three-dimensional and four-dimensional experiments with non-uniform sampling to assign ~85 % of the backbone resonances. The disordered nature of CC1ΔC223 was supported *via* multiple sequence data analysis algorithms and CD measurements (Fig. S3 A, Fig. 3 B). AUC analysis revealed only elongated monomeric forms of the protein in solution (Fig. 3C). Secondary structure propensities were estimated by generating chemical shift indices (CSI). For this purpose, experimental Cα and Cβ chemical shifts were subtracted from the respective random coil values for each amino acid type. The resulting CSI revealed few and rather scattered deviations from random coil values (Figure 3D). Moreover, the uniform and fast dynamics and chemical shift indices of CC1ΔC223 are consistent with a disordered, highly dynamic and monomeric state in solution (Fig. S3, B to D), similar to the members of the Tau/MAP2/MAP4 family.

**Figure 3.**
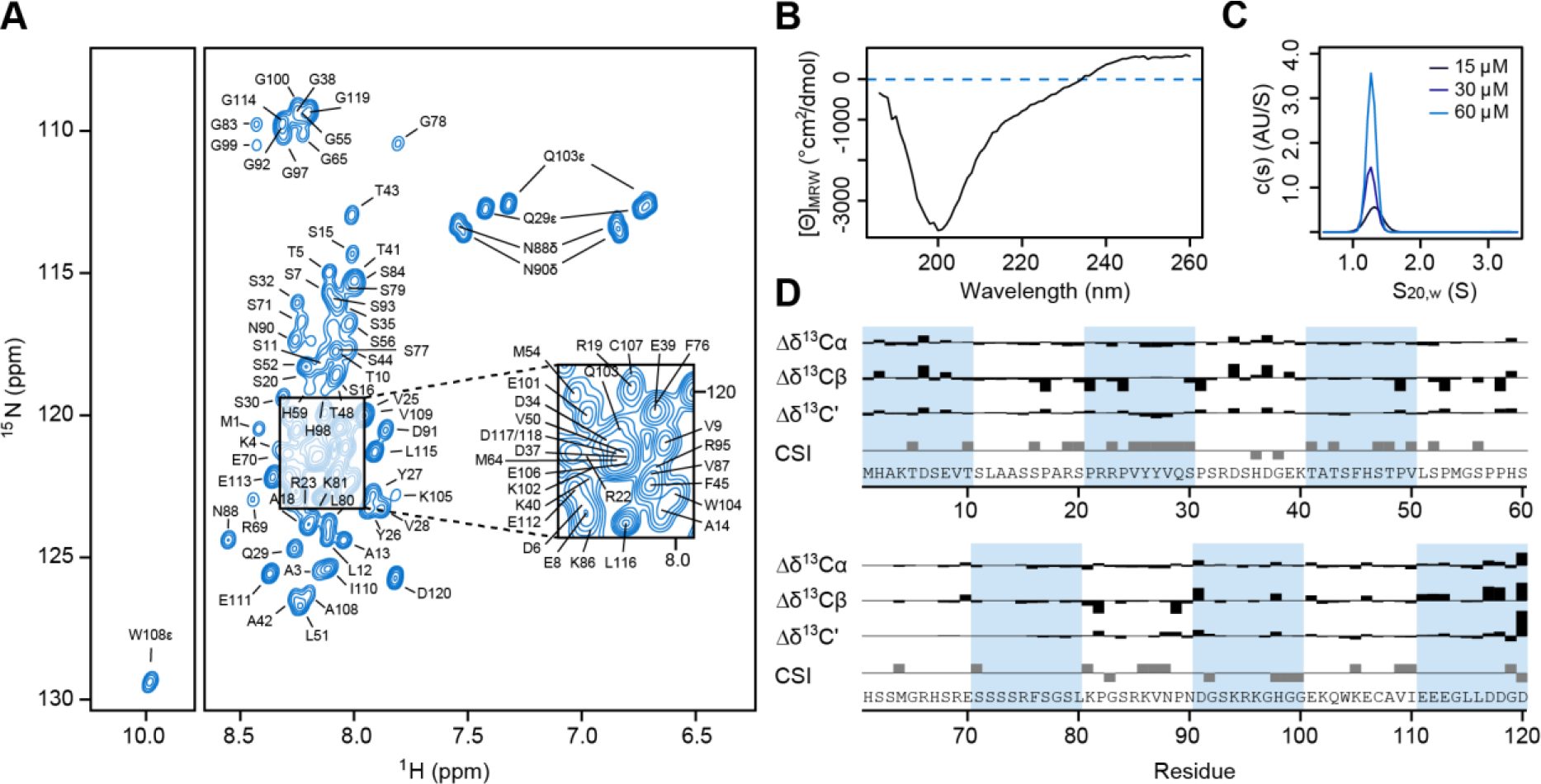
NMR scale protein production and structural characteristics of the CC1 N-terminus. **A.** Assigned ^1^H-^15^N HSQC spectrum of ^15^N-labeled CC1ΔC223 in solution. The low signal dispersion in the ^1^H dimension is characteristic of an intrinsically disordered protein. **B.** Circular dichroism (CD) spectrum of 6xHis-CC1ΔC223 in solution supporting lack of clear structures in the protein. **C.** Analytical ultracentrifugation at three different CC1ΔC223 concentrations showed a single size population at the approximate molecular weight of monomeric CC1ΔC223. The frictional coefficient of 1.7 is characteristic for elongated protein shapes. **D.** ΔδCα, ΔδCβ and ΔδC’ values and chemical shift indices (CSI) for CC1ΔC223 ^36^. Experimental Cα, Cβ and C’ chemical shifts were subtracted from the respective random coil values for each amino acid type.

To study CC1ΔC223-microtubule interactions in a residue-specific manner, we recorded ^1^H-^15^N HSQC spectra of ^15^N-labeled CC1ΔC223 in the presence and absence of paclitaxel-stabilized microtubules. We observed line broadening and vanishing of individual cross-peaks when microtubules were added (Fig. 4, A and B). The effects of the microtubules on the transverse relaxation rate (ΔR_2_) of CC1ΔC223 signals were reversible, independent of the magnetic field, residue specific, did not correlate with the chemical shift changes, and relaxation dispersion experiments did not show contributions of intermediate exchange (Fig. S4, A to G). To conclude, the line broadening is a direct result of CC1ΔC223-microtubule complex formation. Figure 4C shows the intensity ratios of cross-peaks taken from 3D HNCA spectra of ^15^N,^13^C-labeled CC1ΔC223 in the presence and the absence of microtubules (I_*bound*_/I_*free*_) per residue. A significant intensity decrease is observed in four regions, comprising residues ^23^RPVYYVQS^30^, ^45^FHSTPVLSPM^54^, ^74^FSGSLKPG^83^ and ^103^QWKECAVI^110^ (Fig. 4C). Due to signal overlap, the region between residues 60 and 80 is not well covered. We found a clear correlation between the NMR-based microtubule-interaction profile and the hydrophobicity pattern of CC1ΔC223, highlighting the role of hydrophobic interactions (Fig. 4D). The binding motifs are separated by stretches of mobile residues, presumably acting as linkers that are likely to retain a high degree of flexibility thus facilitating a highly dynamic interaction with microtubules. This binding behavior is reminiscent to that of Tau ^23^, and the microtubule-binding regions of CC1ΔC223 also share remarkable similarities in hydrophobicity, size, sequence, and spacing with those of the microtubule-binding regions of Tau(201-320) (Fig. 4E and Fig. S4H).

**Figure 4.**
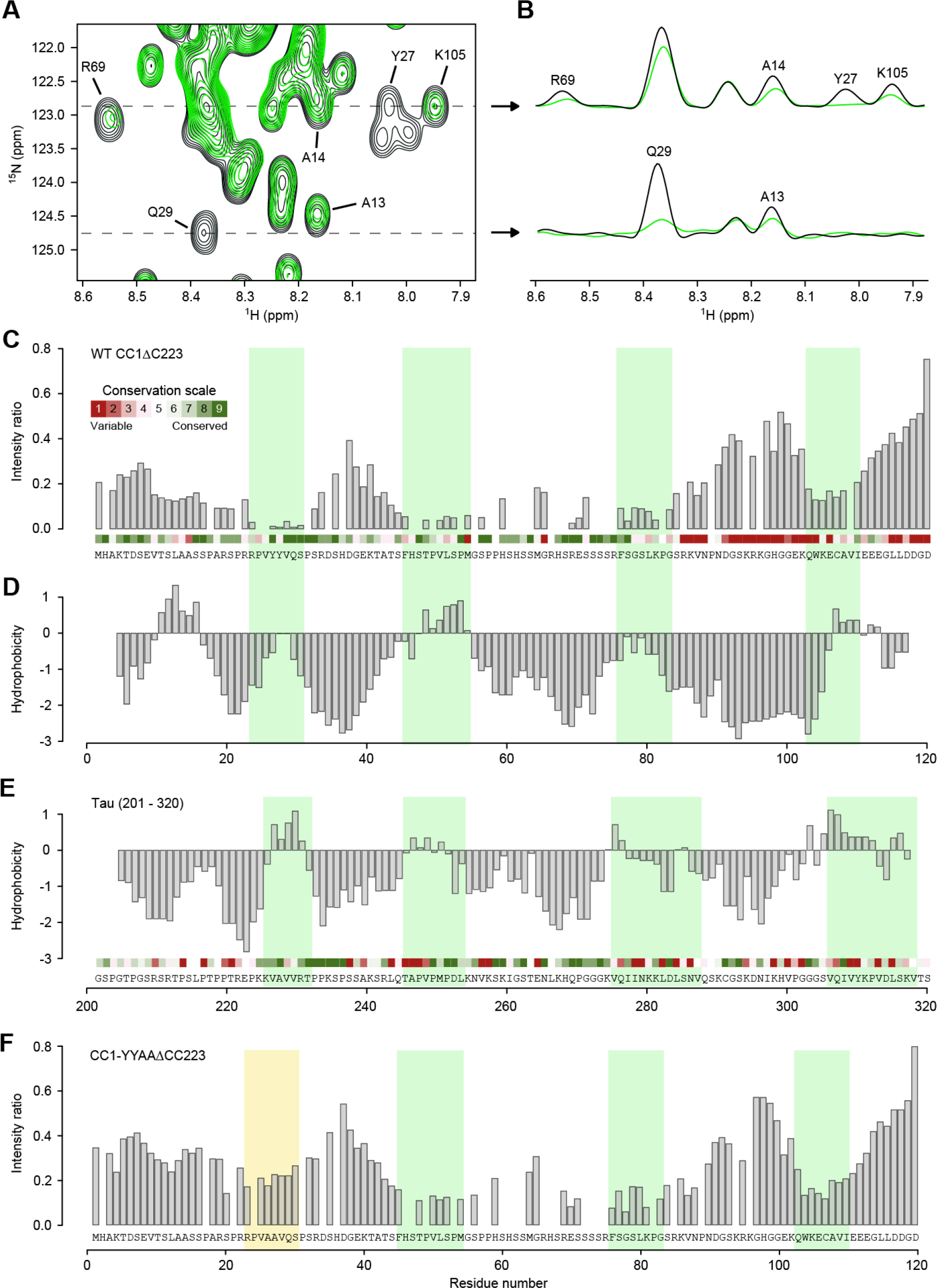
The N-terminus of CC1 binds to paclitaxel-stabilized microtubules via short, hydrophobic and conserved regions. **A.** ^1^H-^15^N HSQC spectrum of free CC1ΔC223 (black) and in the presence of equimolar paclitaxel-stabilized microtubules (green). Selected residues are labeled. **B.** F_2_-cross sections, showing ^1^H-signals, taken along dotted lines in (**A**) at ^15^N frequencies 122.9 and 124.7 ppm. **C.** Intensity ratio of free CC1ΔC223 HNCA signals and in complex with microtubules. Minima are highlighted with green bars. Site-specific evolutionary conservation calculated by CONSURF is plotted above the sequence in a color code (green = conserved, red = unconserved). **D.** Hydrophobicity scores of CC1ΔC223 according to the Kyte-Doolittle scale, calculated in a 5-residue window. **E.** Hydrophobicity scores of Tau(201-320) according to the Kyte-Doolittle scale, calculated in a 5 residue window. Sequence conservation is plotted above the sequence like in (**C**). Green bars highlight the interacting regions of Tau with microtubules as in ^23^. **F.** Intensity ratio of free CC1YYAAΔC223 HNCA signals and in complex with microtubules. Mutated N-terminal region highlighted with yellow bar.

### Two Neighboring Tyrosine Residues in the N-terminus of CC1 Contribute to the Microtubule Binding

Microtubule binding of the four regions individually was investigated by Saturation Transfer Difference (STD) NMR measurements (Fig. S5A). The peptides CC1(16-38), CC1(41-64), CC1(65-85) and a positive control peptide Tau(211-242) yielded strong STD intensities in the amide and aromatic regions of the ^1^H spectrum (Fig. S5, B to E). No significant STD effects were observed for a negative control peptide, CC1(83-103), corresponding to the third poorly conserved linker region, and for the most C-terminal region CC1(100-114) (Fig. S5, F and G). Targeting the N-terminal binding site, the exchange of ^26^YY^27^ to alanine in a CC1YYAA(16-38) peptide resulted in a substantially reduced STD profile, corroborating a contribution of these aromatic rings to the interaction (Fig. S5H). Indeed, the same mutation in CC1ΔC223 resulted in significantly reduced signal broadening of residues in the N-terminal region, while the intensity ratios for the C-terminal part remained similar to the wild-type protein (Fig. 3F). Likewise, the mutated CC1ΔC223 bound to microtubules with a lower affinity compared to the wild-type sequence in microtubule spin down assays (Fig. S5, I and J), corroborating an important function of the two tyrosine residues in microtubule binding.

### Mutations of the Two Microtubule-Interacting Tyrosine Residues in CC1 Impair Microtubule-guided CESA Movement and Ability of Plants to Grow on Salt

To assess how mutations in the two microtubule-binding tyrosine residues affect the function of CC1 *in vivo*, we mutated them to alanine in the full-length CC1 (CC1YYAA), fused it N-terminally with GFP, and transformed it into *Arabidopsis thaliana cc1cc2* mutant plants. The *cc1cc2* mutant seedlings display reduced growth and crystalline cellulose content on salt-containing media ^5^. These phenotypes were not restored in *cc1cc2* GFP-CC1YYAA seedlings when grown on salt-containing media as compared to controls (Fig. 5, A to C) Spinning-disc confocal microscopy showed GFP-CC1YYAA signals as distinct foci at the plasma membrane (Movie S2) and within cytoplasmic compartments in dark-grown Arabidopsis hypocotyl cells, in accordance with reports on GFP-CC1 (^5^; Fig. S6, A to C). GFP-CC1 co-localizes and migrates with tdTomato(tdT)-CESA6, which is an important subunit of the CSC ^24^, at the plasma membrane ^5^. Notably, the GFP-CC1YYAA also co-migrated with tdT-CESA6 at the plasma membrane (Fig. S6, A to E) (Pearson correlation coefficient r = 0.74 ± 0.06; 6 cells from 6 seedlings and 3 biological replicates). However, in contrast to GFP-CC1, the migration of GFP-CC1YYAA was largely independent of cortical microtubules (mCherry (mCh)-TUA5; ^25^; Fig. 5, D to F). This indicates that reduced microtubule binding of GFP-CC1YYAA either directly affects the ability of CSCs to engage with microtubules, or that the microtubule array is mis-regulated and cannot fulfil its guiding function anymore.

**Figure 5.**
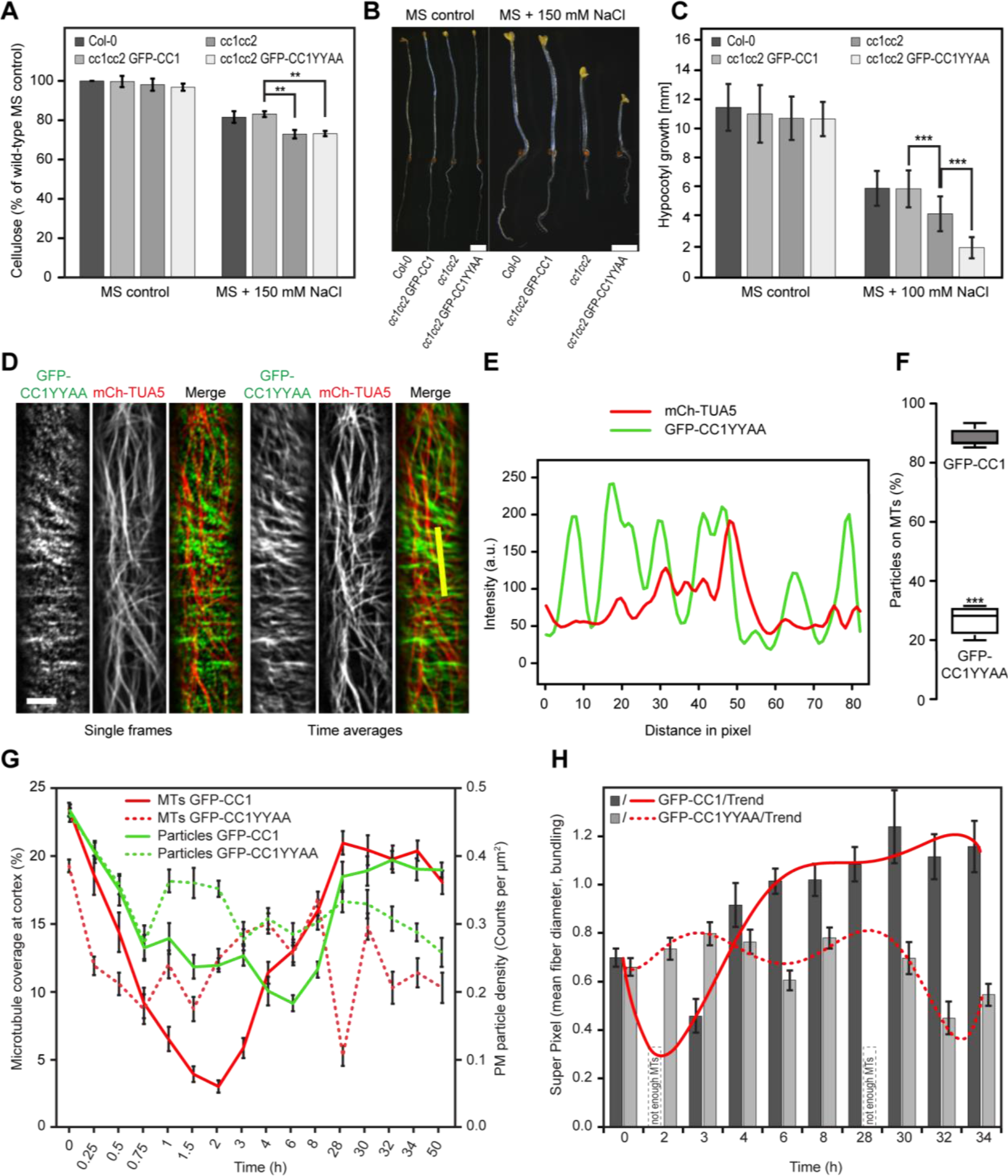
Mutations in the first microtubule binding region of CC1 impair salt tolerance of plants because of mis-regulated microtubule organization upon salt stress. **A.** Cellulose levels in seedlings grown as in (**B**). Values are means +/− SD expressed as % cellulose of wild-type seedlings grown on MS control media. N = 3 biological replicates with 3 technical replicates each. Unpaired *t*-test; ** p-value ≤ 0.01. **B.** Seedlings germinated and grown for two days on MS plates and then transferred to either MS control plates or MS plates supplemented with 150 mM NaCl and grown for additional 5 days. Scale bar = 2 mm. Please be aware that the images were stitched with Leica LAS X Life Science software. **C.** Quantification of hypocotyl elongation of seedlings grown on MS plates for three days and then transferred to either MS control plates or MS plates supplemented with 100 mM NaCl and grown for additional 4 days. Values are mean +/− SD, N = 30 seedlings, 10 seedlings each per three independent experiments. Unpaired *t*-test; *** p-value ≤ 0.001. **D.** GFP-CC1YYAA and mCh-TUA5 in dual-labeled three-day-old *cc1cc2* etiolated hypocotyls (left panels; single frame, right panels; time average projections). Scale bars = 5 μm. **E.** Fluorescence intensity plot of GFP-CC1YYAA and tdT-CESA6 from transect in (**D**) along the depicted yellow line. Note that the GFP signal does not substantially correlate with the mCherry signal. **F.** Quantification of GFP-CC1 and GFP-CC1YYAA fluorescent foci on cortical microtubules in a 50×50 pixel area of five individual time-lapse images, N = 5 cells from 5 seedling and 3 independent experiments (box plots: Center lines show the medians; box limits indicate the 25th and 75th percentiles; whiskers extend to the minimum and maximum). Unpaired *t*-test; *** p-value ≤ 0.001. **G.** Quantification of microtubule and GFP-CC (GFP-CC1 or GFP-CC1YYAA) coverage at the cell cortex and plasma membrane, respectively, after exposure of *cc1cc2* seedlings to 200 mM NaCl as in an experiment shown in figure **S6F**. Time indicates time after salt exposure. Values are mean +/− SEM, n = 27 cells from 3 seedlings per time point and 3 independent experiments. Two-way ANOVA analysis of microtubule coverage; p ≤ 0.001 (genotype), p ≤ p ≤ 0.001 (time), p ≤ 0.001 (genotype x time). Two-way ANOVA analysis of GFP-CC protein density; p ≤ 0.01 (genotype), p ≤ 0.001 (time), p ≤ 0.001 (genotype x time). **H.** Quantification of microtubule bundling after exposure of *cc1cc2* GFP-CC1 /GFP-CC1YYAA seedlings to 200 mM NaCl as in an experiment shown in figure **S6F**. The salt adjusted microtubule array in GFPCC1 seedlings shows increased bundling after exposure to salt while the array GFPCC1YYAA seedlings does not. Values are mean +/− SEM, n = 27 cells from 3 seedlings per time point and 3 independent experiments. Two-way ANOVA analysis of microtubule bundling (excluding T2 and T28); p ≤ 0.001 (genotype), p ≤ 0.001 (time), p ≤ 0.001 (genotype x time).

To investigate if the CC1YYAA can sustain microtubule and CSC function during salt exposure, we exposed seedlings to 200 mM salt and recorded time series of microtubule (mCh-TUA5) and CC1 (GFP-CC1 or GFP-CC1YYAA) behavior (Fig. S6F). The GFP-CC proteins (either GFP-CC1 or GFP-CC1YYAA) were considered as proxy for the CSC behavior because they co-localize and migrate together with tdT-CESA6. In agreement with ^5^, the microtubule array and cellulose synthesis were restored within 28 hours of salt exposure in the GFP-CC1-complemented *cc1cc2* seedlings (Fig. 5G). However, the *cc1cc2* GFP-CC1YYAA-complemented seedlings largely mimicked the *cc1cc2* mutant seedlings and failed to restore the microtubule array and cellulose synthesis during the course of the experiment (Fig. 5G). Interestingly, while the *cc1cc2* GFP-CC1 line showed increased microtubule bundling of the salt-adjusted microtubule array, the *cc1cc2* GFP-CC1YYAA cells failed to do so (Fig. 5F). Furthermore, the microtubule dynamics differed in the GFP-CC1 and GFP-CC1YYAA cell lines (Fig. S6, G and H), indicating that the microtubule dynamics and bundling are key to build a salt-tolerant microtubule array. Hence, the YY-containing region of CC1 is necessary to sustain microtubule array organization and cellulose synthesis during salt stress.

## Discussion

Abiotic stress, such as soil salinity, substantially impacts plant growth ^26^ and thus dramatically compromises global agricultural productivity (~50-80 % loss in yield; ^27,28^). Unravelling molecular mechanisms that can be used to engineer plants for better stress tolerance is therefore of urgent importance. We propose that the microtubule-binding regions of CC1 interact transiently with tubulin heterodimers, promoting polymerization by increasing the local tubulin concentration, stabilizing, and bundling microtubules, which would support the formation of a stress-stable microtubule array. The hydrophobic interactions of the CC1-microtubule complex could permit a more robust binding under conditions of high ionic strength, corroborating the importance of the protein’s function during salt stress. The two tyrosine residues in the most N-terminal microtubule-binding region of CC1 are key to the microtubule binding, both *in vitro* and *in vivo*. Mutations in these residues disrupted microtubule-guided CSC movement and led to failure in the generation of a stress-tolerant microtubule array.

Our results show that CC1ΔC223 functions remarkably similar to Tau. Both are intrinsically disordered proteins that can diffuse bidirectionally along the microtubule lattice ^10,23^. A Tau fragment encompassing the four NMR-derived microtubule binding regions (Tau(208-324); TauF4) joins microtubules wall-to-wall similar to that of CC1ΔC223 ^29^. In-depth NMR studies using TauF4 ^30^, and cryo-EM studies on full-length Tau ^8^, proposed that Tau spans multiple tubulin heterodimers along the microtubule principal axis when bound to microtubules. The equivalence of cross-linked positions on α-tubulin and the longitudinal decoration in the gold-labeling experiments suggest a similar interaction of CC1ΔC223 with microtubules ^18^. The functional and structural analogies between Tau and CC1 are further reflected in the fact that both Tau and CC1 are relevant for the organism to function during stress conditions; CC1 promotes cellulose synthesis during salt stress ^5^, whereas Tau has emerged as a key regulator of stress-induced brain pathology in mice and oxidative stress in cultured fibroblasts ^31,32^. Furthermore, similar to the tyrosine to alanine mutations in CC1, disease-related mutations in Tau cause distinct defects in microtubule organization ^33^. While the typical PGGG-containing repeats of the Tau microtubule-binding domain (R1-R4) are not obvious from the CC1 sequence, the two proteins do contain four similarly spaced hydrophobic microtubule-binding regions (regions 1-4 in Fig. S4H). A sequence comparison of these four regions reveals a surprisingly high number of identical or similar residues Fig. S4H, bottom), implying evolutionary convergence of the microtubule-binding mechanism.

While the microtubule-regulating mechanisms appear to be comparable between Tau and CC1 (Fig. 6), other features of the two proteins are clearly different. For example, Tau is a cytoplasmic protein with an N-terminal projection domain that regulates microtubule spacing ^34^, whereas CC1 contains a putative transmembrane domain and is closely connected to the CSC ^5^. The proteins are thus situated in different cellular contexts that will influence their modes of operation. Notably, the microtubule arrays have different design principles in animal and plant cells. The centrosome-coordinated microtubules in animal cells typically radiate from the cell center towards the periphery, while growing plant cells have a cortical microtubule array, with evenly distributed microtubules along the cell cortex ^2^. We speculate that the differences in the protein domains and topologies of Tau and CC1 coincide with the microtubule array organization. CC1 is a core component of the CSC (Fig. 6), which is primarily localized on cortical microtubule bundles ^3^ and its movement is guided by cortical microtubules in plant interphase cells ^3^. Hence, the CC1 proteins are superbly situated to modulate microtubule dynamics and bundling to optimize cellulose synthesis under different environmental conditions. Our results thus support striking similarities for how plant cells and neurons control microtubule bundling and dynamics in the context of the microtubule array organization. In this setting, the CC1 microtubule-binding motif that contains the two tyrosine residues essential for stress-stable microtubule array formation is most distal to the plasma membrane. Given the local environment of CC1, i.e. being part of the CSC and integral to the plasma membrane, this distal motif might be the most prominently exposed of the four microtubule-binding motifs and therefore also most prominent in the microtubule engagement. Engineering the microtubule-binding properties of this domain, perhaps by design principles of Tau, might improve cellulose synthesis and thus biomass production on salt-affected lands.

**Figure 6.**
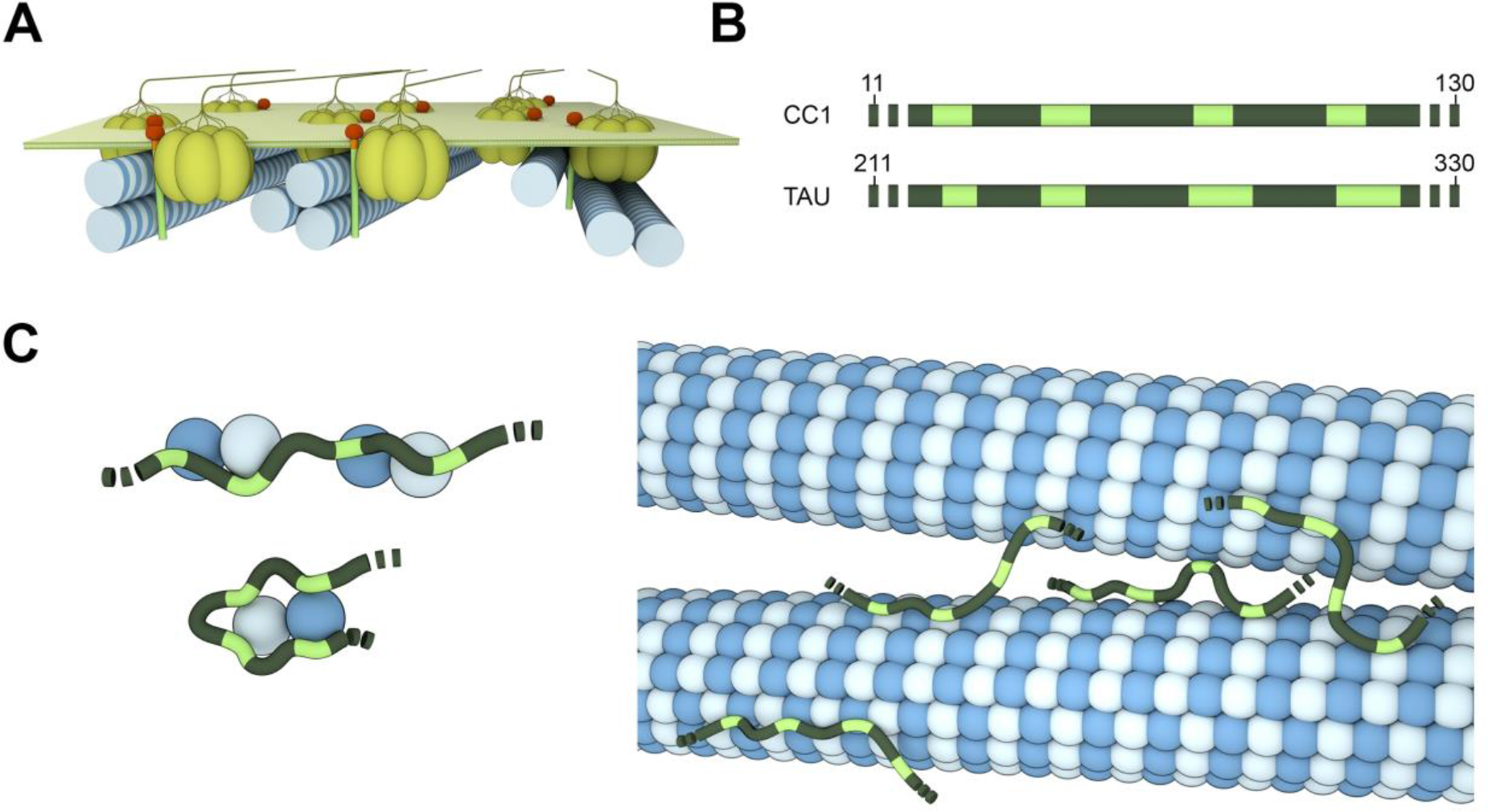
Cartoon Overview of the CC1-microtubule interaction and its similarity to Tau. **A.** CC1 localization in its cellular context as part of the cellulose synthase complex (CSC). CC1 interacts with one or several microtubules while the CSC migrates on cortical microtubules during cellulose production. CC1 regulates microtubule array organization at the same time through its cytosolic N-terminus by e.g. bundling and dynamic modifications. **B.** Microtubule interacting CC1 N-terminus and the corresponding domain of Tau. Both microtubule binding domains show remarkable similarities: four similarly spaced, hydrophobic microtubule binding motifs (highlighted in green) that are spaced by flexible, hydrophilic linker regions. The four hydrophobic sites also share sequence similarities (see Fig. S4H). **C** The dynamic nature of the CC1ΔC223 and Tau binding behaviour suggests that both might be able to bind multiple distinct tubulin dimers *via* their individual binding motifs, thereby increasing the local tubulin concentration, connecting and stabilizing protofilaments or bundling microtubules.

## Acknowledgements

Live cell imaging was performed with equipment maintained by the Center for Microscopy and Image Analysis (University of Zurich) and Scientific Center for Optical and Electron Microscopy (ScopeM, ETH Zurich). We thank Martina Leidert and Natalja Erdmann for help with protein expression and purification. We also thank the Diez group, especially Felix Ruhnow, (B CUBE Center for Molecular Bioengineering, Dresden) for assistance with microtubule imaging and *in vitro* assays. The helper plasmid pBAD-σ32 (I54N) was a gift from Jeffery Kelly ^35^; Addgene plasmid # 59982), pETM11 was kindly supplied by EMBL Protein Production facility (Heidelberg Germany). We thank the Biological Optical Microscopy Platform, the Melbourne Advanced Microscopy Facility, and the Mass Spectrometry and Proteomics Facility (School of Biosciences and Bio21) at the University of Melbourne.

### Funding

S.P was supported by an ARC Discovery grant (DP150103495), a Hermon-Slade Grant (Persson HSF 15/4) and a Future Fellowship grant (FT160100218). J.L.H. was supported by an ARC Future Fellowship (FT130101165). H.E.M. was supported by an ARC DECRA (DE170100054). C.S.R. and C.K. were supported by ETHZ and a SNF grant (2-77212-15). C.K. was supported by a Peter und Traudl Engelhorn-Stiftung fellowship. R.S. received Computational Biology Research Initiative and Early Career Research Grants from the University of Melbourne.

### Author contributions

C.K., A.W., R.S., H.E.M., A.D., B.J.v.R., J.H., C.S.R., H.O. and S.P. designed the research. C.K., A.W., R.S., H.E.M., A.D., M.L., G.A.K., N.C., P.S., F.S. and K.F. performed the research. C.K., A.W., R.S., H.E.M., J.H. and S.P. analyzed data. C.K., A.W., C.S.R., H.O. and S.P. wrote the article. C.K. and A.W. share equal first authorship.

### Competing interests

The authors declare no competing interests.

### Data and materials availability

All data is available in the manuscript or the supplementary materials.

## Supplementary Materials

Materials and Methods

Figure S1 – S6, Table S1

Movies S1 and S2

References (1 –39)

